# Microbiome diversity: A barrier to the environmental spread of antimicrobial resistance?

**DOI:** 10.1101/2023.03.30.534382

**Authors:** Uli Klümper, Giulia Gionchetta, Elisa C. P. Catao, Xavier Bellanger, Irina Dielacher, Peiju Fang, Sonia Galazka, Agata Goryluk-Salmonowicz, David Kneis, Uchechi Okoroafor, Elena Radu, Mateusz Szadziul, Edina Szekeres, Adela Teban-Man, Cristian Coman, Norbert Kreuzinger, Magdalena Popowska, Julia Vierheilig, Fiona Walsh, Markus Woegerbauer, Helmut Bürgmann, Christophe Merlin, Thomas U. Berendonk

**Affiliations:** Technische Universität Dresden, Institute for Hydrobiology, Zellescher Weg 40, 01217 Dresden, Germany; Eawag, Swiss Federal Institute of Aquatic Science and Technology, Department of Surface Waters – Research and Management, 6047 Kastanienbaum, Switzerland; Université de Lorraine, CNRS, LCPME, UMR 7564, Villers-lès-Nancy, France; Université de Toulon, MAPIEM, CS 60584, Toulon, France; Institute of Water Quality and Resource Management, TU Wien, Karlsplatz 13/2261, 1040 Vienna, Austria; Department for Integrative Risk Assessment, Division for Risk Assessment, Data and Statistics, AGES – Austrian Agency for Health and Food Safety, Spargelfeldstraße 191, 1220 Vienna, Austria; University of Warsaw, Faculty of Biology, Institute of Microbiology, Department of Bacterial Physiology, Miecznikowa 1, 02-096 Warsaw, Poland; Department of Biology, Kathleen Lonsdale Institute for Human Health, Maynooth University, Maynooth, Co. Kildare, Ireland; Institute of Virology Stefan S. Nicolau, Romanian Academy of Science, 285 Mihai Bravu Avenue, 030304, Bucharest, Romania; NIRDBS, Institute of Biological Research Cluj-Napoca, 400015, Cluj-Napoca, Romania; Interuniversity Cooperation Centre Water & Health, Austria

**Author notes:** Corresponding author <u>Corresponding author address:</u> Prof. Thomas Berendonk (ORCID: 0000-0002-9301-1803) Technische Universität Dresden Institute of Hydrobiology Zellescher Weg 40 01217 Dresden Germany Mail. Contributed equally to this work (Uli Klümper, Giulia Gionchetta, Elisa C. P. Catao).

**Keywords:** Biodiversity, Microbial barrier, Microbial Invasion, Riverbed microbiome, Soil microbiome, ARG

## Abstract

**Background:** In the environment, microbial communities are constantly exposed to invasion by antimicrobial resistant bacteria (ARB) and their associated antimicrobial resistance genes (ARGs) that were enriched in the anthroposphere. A successful invader has to overcome the biotic resilience of the habitat, which is more difficult with increasing biodiversity. The capacity to exploit resources in a given habitat is enhanced when communities exhibit greater diversity, reducing opportunities for invaders, leading to a lower persistence. In the context of antimicrobial resistance (AMR) dissemination, exogenous ARB reaching a natural community may persist longer if the biodiversity of the autochthonous community is low, increasing the chance of ARGs to transfer to community members. Reciprocally, high microbial diversity could serve as a natural long-term barrier towards invasion by ARB and ARGs.

**Results:** To test this hypothesis, a sampling campaign across seven European countries was carried out to obtain 172 environmental samples from sites with low anthropogenic impact. Samples were collected from contrasting environments: stationary structured forest soils, or dynamic river biofilms and sediments. Microbial diversity and relative abundance of 27 ARGs and 5 mobile genetic element marker genes were determined. In soils, higher diversity, evenness and richness were all significantly negatively correlated with the relative abundance of the majority (>85%) of ARGs. Furthermore, the number of detected ARGs per sample was inversely correlated with diversity. However, no such effects were found for the more dynamic, regularly mixed rivers. Conclusions: In conclusion, we demonstrate that diversity can serve as barrier towards AMR dissemination in the environment. This effect is mainly observed in stationary, structured environments, where long-term, diversity-based resilience against invasion can evolve. Such barrier effects can in the future be exploited to limit the environmental proliferation of AMR.

## Background

The spread of antibiotic resistance genes (ARGs) represents one of the biggest challenges to future human and animal health [1,2]. It has become clear that tackling the issues brought on by the expansion of antimicrobial resistance (AMR) requires a global effort that spans the fields of human, veterinary, and environmental health [3,4]. This perspective, also known as the “One Health” approach, is frequently used as a framework for worldwide efforts to contain the spread of AMR. However, integrating the environmental sphere has, due to its high complexity, proven particularly challenging. Understanding the existing ecological dispersal barriers is necessary to restrict the propagation of AMR in the environment [5,6].

ARGs are ancient, naturally occurring in environmental bacteria and evolved over billions of years [7]. However, during the last century, environmental microbial communities have been constantly subjected to invasion events by antibiotic resistant bacteria (ARB), and their associated ARGs, which have been enriched or released through anthropogenic activities. For example, release and reuse of wastewater effluents or the application of manure to soils are known to be primary conduits of AMR spread in the aquatic and terrestrial environments [8–11]. But even habitats with no direct anthropogenic impact are regularly exposed to lower frequencies of such invasion events, for example, through wildlife and aerial depositions. These natural diffuse sources of AMR-pollution promote the widespread dispersal of AMR over time at low levels [12–16].

Invasion has been defined, both on the micro- and the macro-biological scale, as a process consisting of a sequence of successive steps, namely, (i) introduction, (ii) establishment, (iii) growth and spread, and (iv) impact [17]. For it to be successful, the invader has to overcome the “biotic resistance” towards invasion of the habitat [18]. In theory, this process becomes more difficult with increasing biodiversity [19]. The capacity to exploit the resources provided by a habitat is enhanced when communities exhibit a greater diversity, which in turn reduces opportunities for invaders, hence lowering their persistence [20]. If an invader is phylogenetically, thus presumably also physiologically, close to established community members occupying its specific niche, the invasion process has been shown to become merely stochastic [21]. In contrast, even small-scale disturbance events can considerably increase invasion success by affecting the niche occupancy of resident species [22]. However, such effects strongly depend on the niche partitioning within the given environment rather than diversity alone [23], and might hence be far more pronounced in structured environments with a long-term established niche occupation.

In the context of AMR dissemination, it is reasonable to assume that - as any invader - ARB reaching a natural community may persist longer when the biodiversity of the autochthonous community is low, as this usually coincides with a lower rate of niche occupation. This could result in a longer-term establishment of the ARB invaders, and thus increase the relative abundance of ARGs. Even when the invasion process is not successful in the long term, a slightly prolonged residence time of the invader could have a significant impact on the likelihood of ARGs being horizontally transferred to the endemic microbiota [24]. Alternatively, future invasion events could be favored by reducing the community resilience [25]. Reciprocally, high microbial diversity could impede the spread of ARB and ARGs, thus serving as a potential natural barrier.

Diversity as a limiting factor for AMR invasion was demonstrated in the short-term for laboratory soil microcosms inoculated at different diversities using an artificial dilution-to-extinction approach. Less diverse soil microcosms displayed a far higher likelihood of being invaded by ARBs [26]. Regarding the aquatic dimension, increased invasion success of a resistant *E. coli* strain into river biofilm communities was shown to coincide with a loss in diversity of these communities under stress conditions [27]. Further, diverse microbial communities with a high degree of functional niche coverage, such as activated sludge, have been suggested to provide natural barriers for the proliferation of AMR [28]. However, data suggests that activated sludge communities have also incorporated a particularly high diversity of ARGs encoded on mobile genetic elements (MGEs) [29].

Based on this theoretical and experimental knowledge, we hypothesize that the invasion success and, in consequence, the pervasiveness of AMR into environmental microbiomes is inversely correlated to the diversity of the communities in question. To our knowledge, no prior field-based study has explored whether AMR dissemination is in the long-term indeed related to microbial diversity in terrestrial or aquatic environmental microbiomes. To this end, low impacted environmental samples with no direct anthropogenic depositions were collected. Their ARG levels, beyond natural background levels, are hypothesized to result from the accumulation of invasion success over time from invaders introduced through the previously mentioned low level dispersion routes. Therefore, ARG levels would rise, if after a successful introduction, these invaders were able to either establish themselves long-term in the indigenous microbial communities or transferred their mobile ARG load. Consequently, 172 of such low anthropogenic impact environmental samples were collected during fall/winter 2020/21 across seven European countries. Half of these were taken from forest soils, representing a stationary, structured environment, while the other half was obtained from river sediments and biofilms representing a more dynamic environment. For each sample, the microbial diversity was assessed through bacterial 16S rRNA gene-based amplicon sequencing, while the abundance of 27 AMR markers and 5 MGEs was determined through high-throughput chip-based qPCR to ultimately determine how microbial diversity as a potential barrier determines ARG abundance in low-impact environments.

## Methods

### Soil sampling and processing

The terrestrial campaign consisted of collecting 74 forest soil samples from the seven countries during fall 2020 (Fig. S1). The aim was to obtain sample sets that included samples across a gradient of high and low microbial diversity that are of relatively low anthropogenic impact (Table S1).

From each forest location five single core samples (Pürckhauer drill, Buerkle™, Germany) were extracted from a depth of 0-25 cm along two 10 m virtual diagonals laid across the sampling location in the form of an X-pattern. 200 g of each of these five subsamples were combined in an aseptic plastic bag, thoroughly homogenized and transferred to the laboratory at 10 °C. From the composite sample aliquots of 20 g were sieved (2 mm mesh size) and stored at -20 °C. DNA extraction was performed using the DNeasy PowerSoil Pro Kit (Qiagen, Germany) according to the manufacturer’s instructions. To obtain DNA from a total of 1 g of each sieved soil sample four replicates of 0.25 g each were extracted in parallel and combined thereafter. The quality and quantity of the extracted DNA was assessed spectrophotometrically.

### Riverbed material sampling and processing

The aquatic campaign included the collection of 98 river samples from seven countries (Austria, France, Germany, Ireland, Poland, Romania, and Switzerland) during the fall/winter 2020/21 (Fig. S1). The locations were selected to obtain samples across a gradient of high and low microbial diversity that are of relatively low anthropogenic impact (e.g., not exposed to wastewater treatment plant effluents) (Table S2). At each site, the substrate best representing sessile, non-phototrophic, oxygenated microbial communities in the chosen riverbeds, was sampled. Either epilithic biofilms from the undersides of rocks to avoid phototrophic communities for those streams dominated by rock/gravel, or oxygenated sediment for streams dominated by fine sediment were collected. Specifically, for epilithic biofilm samples, five individual rocks, collected from a shadowed sample area from a riverbed length of approximately 10 meters, were gently scraped from the bottom surface using a sterile toothbrush, and combined to create a composite river biofilm sample. Repeated rinsing with sterile water in a 50 mL falcon tube was performed to collect the biomass. If no rocks or rock biofilms were available, fine surface sediment from shaded areas was sampled. In this case, the upper layer (∼ 5 cm) of sediment was collected using a 50 mL falcon tube. Five sediment cores were combined at equal weight to obtain one composite sample.

All collected samples were gently homogenized and transported to the laboratory on ice. Then, samples were centrifuged (4,000 rpm for 5 min at 4 °C), the supernatant removed, pellets weighted and stored at -20 °C. DNA extraction was performed using the DNeasy PowerSoil Pro Kit (Qiagen, Germany) according to the manufacturer’s instructions. Extraction blanks were used to confirm the absence of DNA contamination. The quality and quantity of extracted DNA was assessed spectrophotometrically.

### Amplicon sequencing and analyses of sequence datasets

To analyze the microbial diversity and taxonomic composition of the samples, DNA extracts were sent to the IKMB sequencing facility (minimum 10,000 reads per sample; Kiel University, Germany). Illumina MiSeq amplicon sequencing of the bacterial 16S rRNA gene was performed using primers targeting the V3-V4 region (V3F: 5′-CCTACGGGAGGCAGCAG-3′ V4R: 5′-GGACTACHVGGGTWTCTAAT-3) [30]. Sequences were collectively analyzed using DADA2 [31] in QIIME2 [32]. Forward and reverse reads were merged into amplicon-sequence variants (ASV), at 99% sequence similarity. Prior to downstream analyses, the *filter-features* and *filter-table* scripts were applied in QIIME2 to clean ASV and taxa tables by removing unclassified and rare ASVs with a frequency of less than 0.1% of the mean sample depth. The corresponding river and soil ASV tables consisted of 15,473 ASVs (river dataset) and 14,464 ASVs (soil dataset). All sequencing data was submitted to the NCBI sequencing read archive under project accession number PRJNA948643.

### High-throughput qPCR of ARGs and genetic markers for MGEs

To determine the relative abundance of target genes in each sample, DNA extracts were sent to Resistomap Oy (Helsinki, Finland) for HT-qPCR analysis using a SmartChip Real-time PCR system (TaKaRa Bio, Japan). The target genes included 27 ARGs and 5 MGEs (Table S3) [33]. In addition, the 16S rRNA gene and the anthropogenic fecal pollution indicator crAssphage [34,35] were quantified. The protocol was as follows: PCR reaction mixture (100 nL) was prepared using SmartChip TB Green Gene Expression Master Mix (TaKara Bio, Japan), nuclease-free PCR-grade water, 300 nM of each primer, and 2 ng/μL DNA template. After initial denaturation at 95 °C for 10 min, PCR comprised 40 cycles of 95 °C for 30 s and 60 °C for 30 s, followed by melting curve analysis for each primer set. A cycle threshold (CT) of 31 was selected as the detection limit [36,37]. Amplicons with non-specific melting curves or multiple peaks were excluded. The relative abundances of the detected gene to 16S rRNA gene were estimated using the ΔCT method based on mean CTs of three technical replicates [38].

### Data analysis and visualization

To characterize the alpha diversity of the terrestrial and aquatic microbial communities, the diversity indices Chao1 richness, Shannon diversity and Pielou evenness were calculated using the *core-metrics-phylogenetic* script in QIIME2 [32] on rarefied data, with samples with an insufficient sampling depth to reliably assess diversity removed from the datasets. The taxonomy of these microbial communities was classified based on the SILVA classifier (version 138, [39]). Beta-diversity dissimilarities among microbial communities were assessed using a principal coordinate analysis (PCoA) based on Bray-Curtis distance matrices [40].

Differences in the relative ARG abundance for aquatic and terrestrial samples were transformed to a log10 scale, and displayed using the package *ComplexHeatmap* [41]. To explore the degree of correlation present among the ARGs in both environments, pairwise correlations analysis based on Spearman rank were assessed. Only correlations showing coefficients |ρ| > 0.75 were considered significant at p < 0.05 after Benjamini-Hochberg correction for multiple testing. The connectivity in relative abundance among ARGs in aquatic and terrestrial samples was displayed through network analysis, performed using the *picante* package [42] based on significant Spearman correlations. The network visualization was generated using the open-source software Gephi v8.2. To test the correlation between resistome and bacterial diversity, correlation analysis between relative ARG abundance and the calculated alpha diversity metrics was performed based on Spearman rank correlation or Pearson correlation followed by Bonferroni correction for multiple testing. Individual groups of data were compared using One-Way ANOVA with post-hoc Tukey HSD tests. Significant differences of correlations from 0 were tested using a t-test. All statistics and plots were produced using R version 4.2.0 [43].

## Results

### Assessing diversity in the river and soil dataset

Two complementary sets of samples were obtained from a total of 94 river as well as 70 soil samples. When assessing the beta diversity of the microbial communities, soil samples differed significantly and with a large effect size from those obtained from river biofilms (PERMANOVA, pseudo-F = 14.73, p < 0.001, n = 148) and river sediments (pseudo-F = 5.53, p < 0.001, n = 78) (Fig. 1A). No clear distinction was observed comparing sediments and epilithic biofilms (pseudo-F = 2.77, n = 94). Consequently, these samples were subsequently grouped to create the combined river dataset. No clear trends with regards to the countries of origin were observed for either dataset, but the spread of the entire dataset exceeded the spread of any sample set from a single country and some river communities from Romania formed a distinct cluster (Fig. 1A).

**Figure 1.**
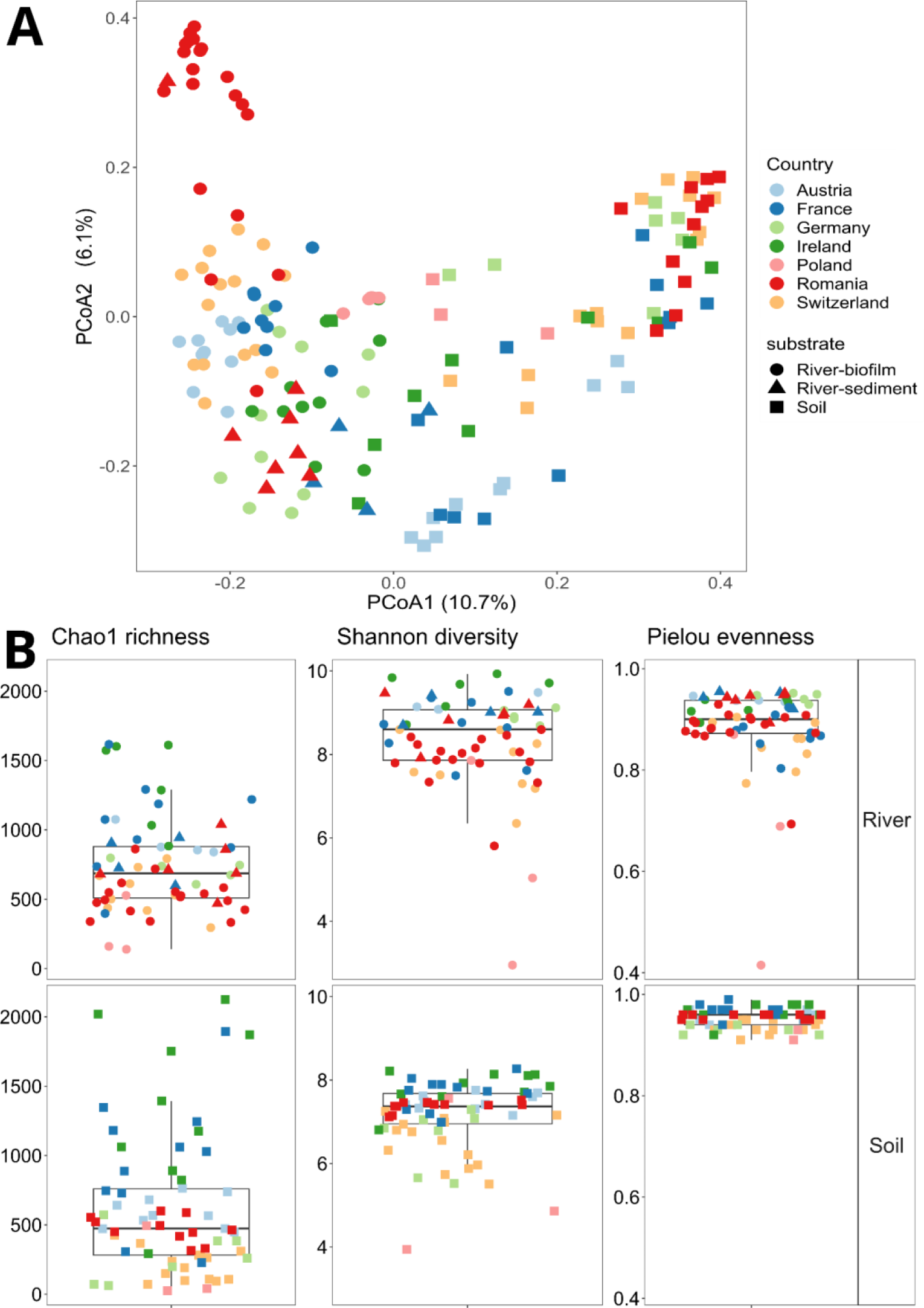
Diversity of the river and soil datasets. Symbols depict sample type, colors code for the country of origin. A) PCoA of the beta diversity based on Bray Curtis distance of ASV relative abundance data from riverbed materials (sediments and biofilms) and soil. B) Alpha-diversity indices (Chao1 richness, Shannon diversity and Pielou evenness) from riverbed materials (top) and soil (bottom) collected from the seven countries.

The aquatic samples contained 19 phyla with an average relative abundance above 1% and were, throughout, dominated by bacteria belonging to the phyla *Proteobacteria*, *Bacteroidota* and *Actinobacteriota* (0.26 ± 0.09, 0.17 ± 0.13, 0.12 ± 0.09) (Fig. S2). In the soil dataset *Acidobacteria*, *Actinobacteriota* and *Proteobacteria* (0.18 ± 0.04, 0.15 ± 0.02, 0.13 ± 0.03) dominated (Fig. S3). No significant differences in dominant phyla between samples based on country of origin were observed. CrAssphage, a common indicator for recent anthropogenic fecal pollution, was undetected in the entirety of soil samples and in 78% of the river samples. For the latter it remained at low relative abundance below 10^-5^ copies per copy of the bacterial 16S rRNA gene, confirming that samples were indeed of low anthropogenic impact origin. For the three main alpha-diversity metrics - Chao1 richness, Shannon diversity and Pielou evenness - high and low biodiversity samples for each of the two datasets, rivers and soils, were obtained (Fig. 1B). The main distinction between the two datasets was the significantly higher level of Pielou evenness in the dataset from the structurally stable soil (0.95 ± 0.02) compared to the dynamic river environment (0.89 ± 0.08) (*p* < 0.0001, *f* = 48.78, One-Way ANOVA). The differences obtained subsequently allowed these diversity metrics to be used as test variables for correlation with ARG abundance.

### Resistome diversity and abundances

In both resistome datasets, the *aac(3)-VI* gene conferring aminoglycoside-resistance was the most abundant (Fig. 2). In the river dataset, genetic determinants for sulfonamide (*sul1*), vancomycin (*vanA*), colistin (*mcr1*) and phenicol (*floR*) resistance clustered together as dominant, followed by other ARGs that promote resistance to macrolides, lincosamides and streptogramins B - (*mphA*), β-Lactam (*blaCTX-M2, blaCTX-M1, blaCMY2*), phenicol (*cmlA*) and aminoglycoside (*aac(6’)- lb3,* a*ph(3’)-Ib*) antibiotic classes. ARGs conferring resistance to quinolone, trimethoprim and tetracycline were less abundant, although in certain soils, particularly from Poland and Romania, the corresponding abundance of individual genes (e.g., *tet(W), qnrS, drfA*) was higher (Fig. 2A). In general, considerable clustering of samples was observed. The Irish samples clustered together with a few Romanian and one German sample separately from the rest, and displayed an overall lower relative ARG abundance with the exception of the *blaTEM* gene, which displayed a particularly high abundance (Fig. 2A).

**Figure 2.**
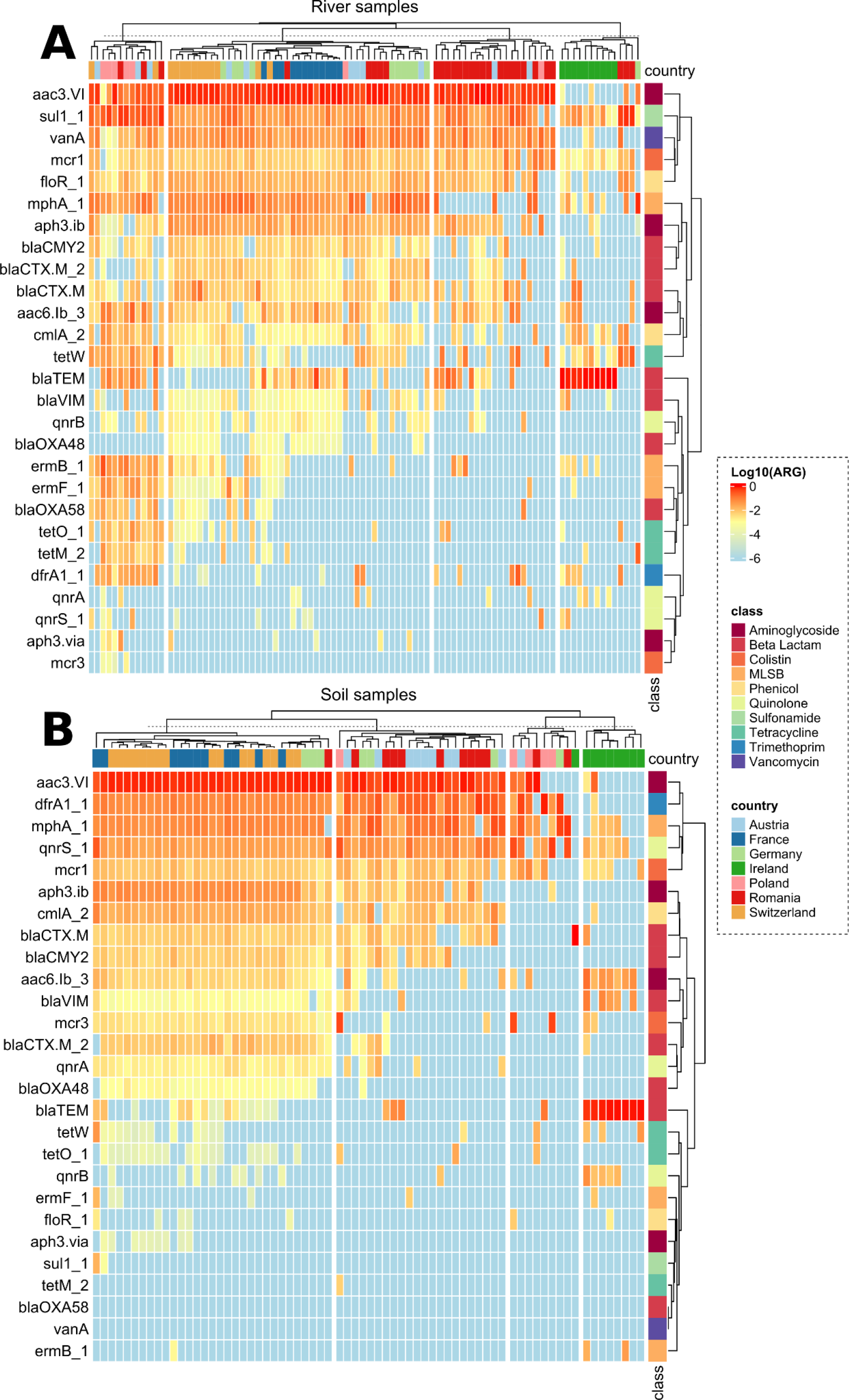
Heatmap of relative ARG abundances in the river (A) and soil (B) dataset. Values are displayed after transformation to log10 scale. The list of ARGs is presented based on similarity in abundance patterns and displayed from high abundance (red) to below the detection limit (blue). Color coding on the right displays the class of antibiotics they confer resistance to. Samples are ordered according to similarity in ARG profiles and color coded based on country of origin displayed on top.

In the soil sample set, the *acc(3)-VI* (aminoglycoside), *qnrS1* (quinolone), *mphA* (MLSB) and *dfrA1* (trimethoprim) genes clustered together as the most abundant determinants in most countries (Fig. 2B). ARGs conferring resistance to colistin (*mcr-1*), Phenicol (*cmlA2*), β-Lactams (*blaCMY-2, blaCTX-M*) and aminoglycoside (a*ph(3’)-Ib, aac(6’)-lb3*) were detected in most countries at intermediate abundances, but with highest values in Switzerland and France. The colistin ARG *mcr-3* was the sole ARG below the limit of detection for all samples, therefore it was not used for any further analysis. Accordingly, the Swiss and French soil resistomes clustered together as most of the ARGs were found at higher and similar abundances compared to the other countries (Fig. 2B). Irish soils displayed again the lowest number of ARGs detected and similar to the river dataset the abundance of *blaTEM* was significantly elevated in Ireland compared to other countries (Fig. 2B).

### Higher degree of correlation between relative ARG abundances in soils compared to rivers

In both datasets, a high number of significant correlations were found between the relative abundance of different ARGs. Out of all potential comparisons between individual relative ARG abundances 140 out of 351 (27.40%) for rivers and 144 out of 325 (44.31%) for soils were significantly positively correlated with each other (p < 0.05, Spearman) (Fig. 3 A,B). Only in the case of aminoglycosides resistance gene *aph(3’)-Ib* and trimethoprim resistance gene *dfrA1* in the river dataset, a single negative correlation in relative abundance could be detected. While in the river environment the distribution of correlations did not follow a clear pattern, in soil a distinct cluster of 17 ARGs that are highly correlated among each other was observed. This cluster included ARGs conferring resistance to antibiotics belonging to different classes, namely vancomycin, tetracyclines, quinolones, polymyxins, phenicols, MLSB, β-Lactams and aminoglycosides. Further the average correlation coefficient of the significant comparisons (Rs) among all soil ARGs was significantly higher (Rs = 0.40 ± 0.33) than among the river dataset (Rs = 0.30 ± 0.29; p < 0.0001, ANOVA) (Fig. 3). This increased level and higher degree of consistency of correlations between ARGs in soil was further confirmed using network analysis, where the average degree of connections of each ARG with the remaining ones was significantly higher in soil (10.533) compared to the river dataset (5.692) (Fig. S4).

**Figure 3.**
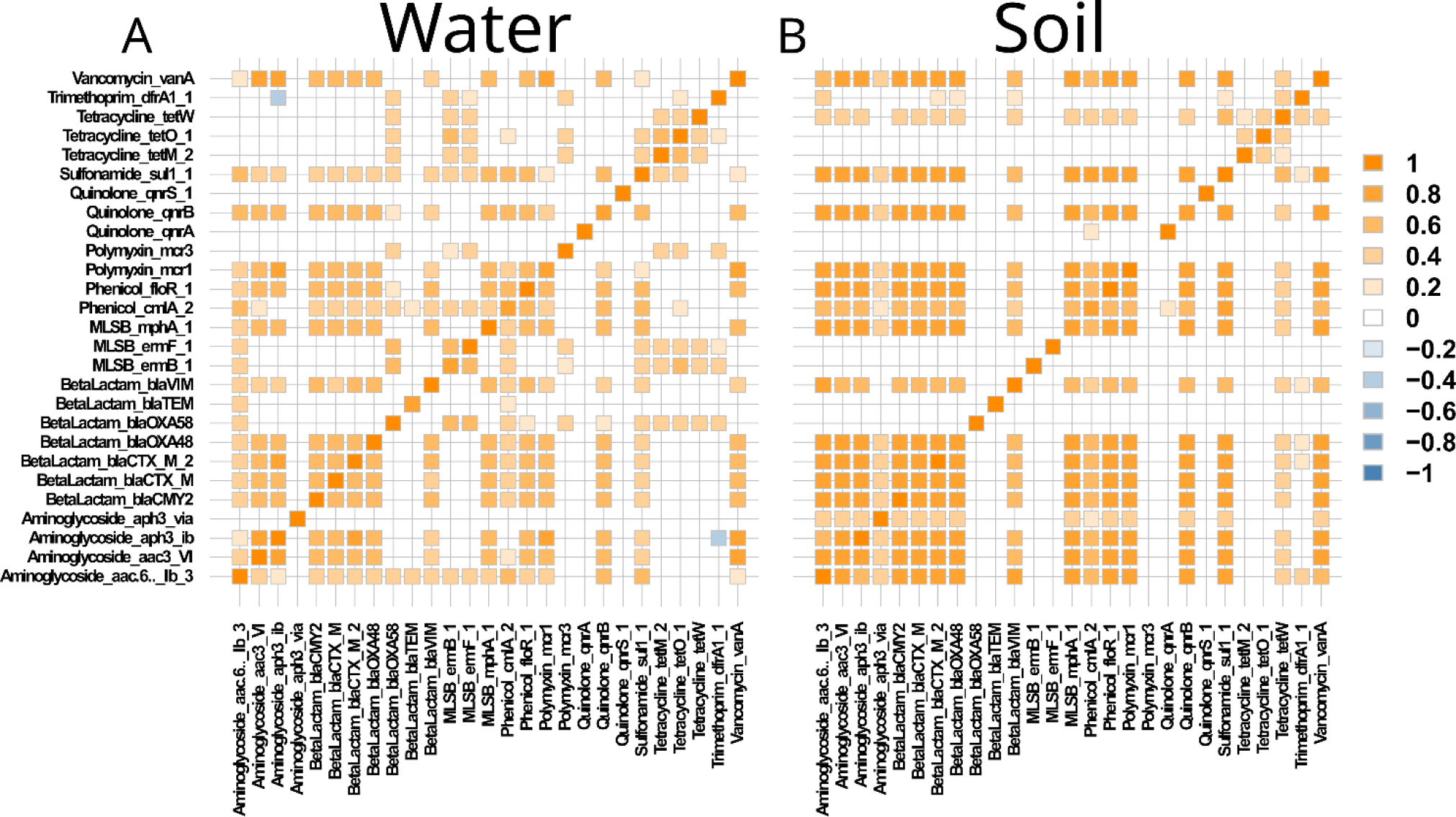
Correlation analysis between relative ARG abundances: Pairwise correlations in river (A) and soil (B) microbial communities based on Spearman rank correlation. Only significant comparisons (p < 0.05 after Benjamini-Hochberg correction for multiple testing) are shown. Intensity of colors displays strength of correlation with orange depicting positive and blue depicting negative correlation.

No significant correlation of the observed ARG number or the relative abundance of any ARGs with the relative abundance of crAssphage was obtained for the river dataset (all p > 0.05, Spearman) while crAssphage was absent in the soil dataset. Thus, it again demonstrates that results are not directly impacted by recent anthropogenic fecal pollution.

### Diversity as a barrier to ARG spread

To determine whether higher diversity could lower the long-term invasion success of ARGs by the communities, we examined correlations between the diversity metrics and the number of detected ARGs as well as the relative abundance of each individual gene in both datasets. On average, 18.44 ± 5.61 of the 27 ARGs tested were successfully detected in samples from the river dataset. No clear trend could be observed in correlation between the number of detected ARGs and any of the three diversity metrics. While all three correlations were negative, none were statistically significant (rPielou = -0.02, rShannon = -0.04, rChao1 = -0.11, all p > 0.05; Pearson correlation; Fig. 4 A-C).

**Figure 4.**
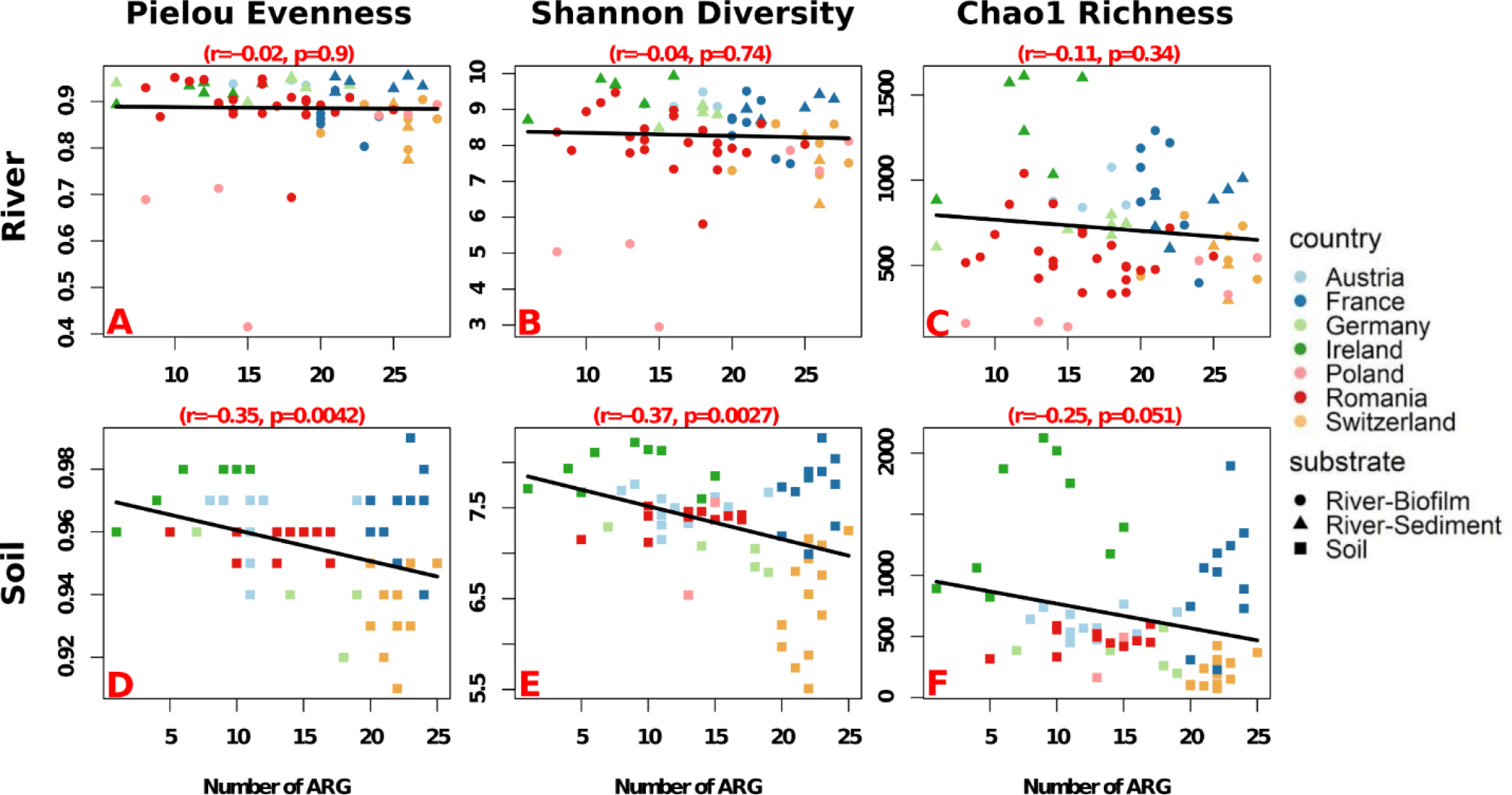
Correlation analysis of the number of ARGs detected per sample with diversity metrics based on Pearson correlation with Bonferroni correction for multiple testing. Linear correlations from river environmental samples with Pielou Evenness (A), Shannon Diversity (B) and Chao1 Richness (C). Linear correlations from soil environmental samples with Pielou Evenness (D), Shannon Diversity (E) and Chao1 Richness (F). Colors depict the country of sample origin and the symbols depict the sample type.

Slightly, but significantly, less ARGs per sample (15.95 ± 6.05; p = 0.014, *f* = 6.11, One-Way ANOVA) were successfully detected in the soil dataset. Contrary to the river dataset, higher diversity in soils correlated with a lower number of detected ARGs. This negative correlation was significant based on Spearman rank correlation analysis for Pielou evenness (r = -0.35, p = 0.0042) and Shannon diversity (r = -0.37, p = 0.0027) (Fig. 4 D,E). Similarly, for Chao1 Richness an inverse correlation with the number of ARGs detected was observed, however, barely not significant (r = -0.25, p = 0.051) (Fig. 4 F). These results provided a first indication that diversity-based barrier effects might indeed exist, at least in the more structured soil environment. Although diversity had a significant impact, the effect sizes of 25-37% suggest that, as expected for complex environmental datasets, diversity is only one of multiple interacting drivers of the observed trends. To further test this hypothesis the relative abundance of each individual ARG was correlated with the obtained diversity metrics. To account for zero-inflation during correlation, only those ARGs that were found in at least 25% of the samples of the respective dataset were tested. In the river dataset, similar to the number of ARGs no clear trends were observed for the relative abundance of any of the tested ARGs (Fig. 5 A-C). The only correlation considered significant based on Spearman rank correlation (with Bonferroni correction for multiple testing) was a positive one between the *blaTEM* gene and the Chao1 richness (RS = 0.42, p = 0.0003). For the remaining combinations of ARGs and diversity metrics only slight negative or slight positive correlation trends (RS = -0.24 - 0.33, all p > 0.05) were observed. Among those non-significant trends, no obvious patterns emerged. In fact, the average Spearman’s *rho* of the tested ARGs for each of the three diversity metrics was near 0 (p > 0.05; t-test with Bonferroni correction for multiple testing).

**Figure 5.**
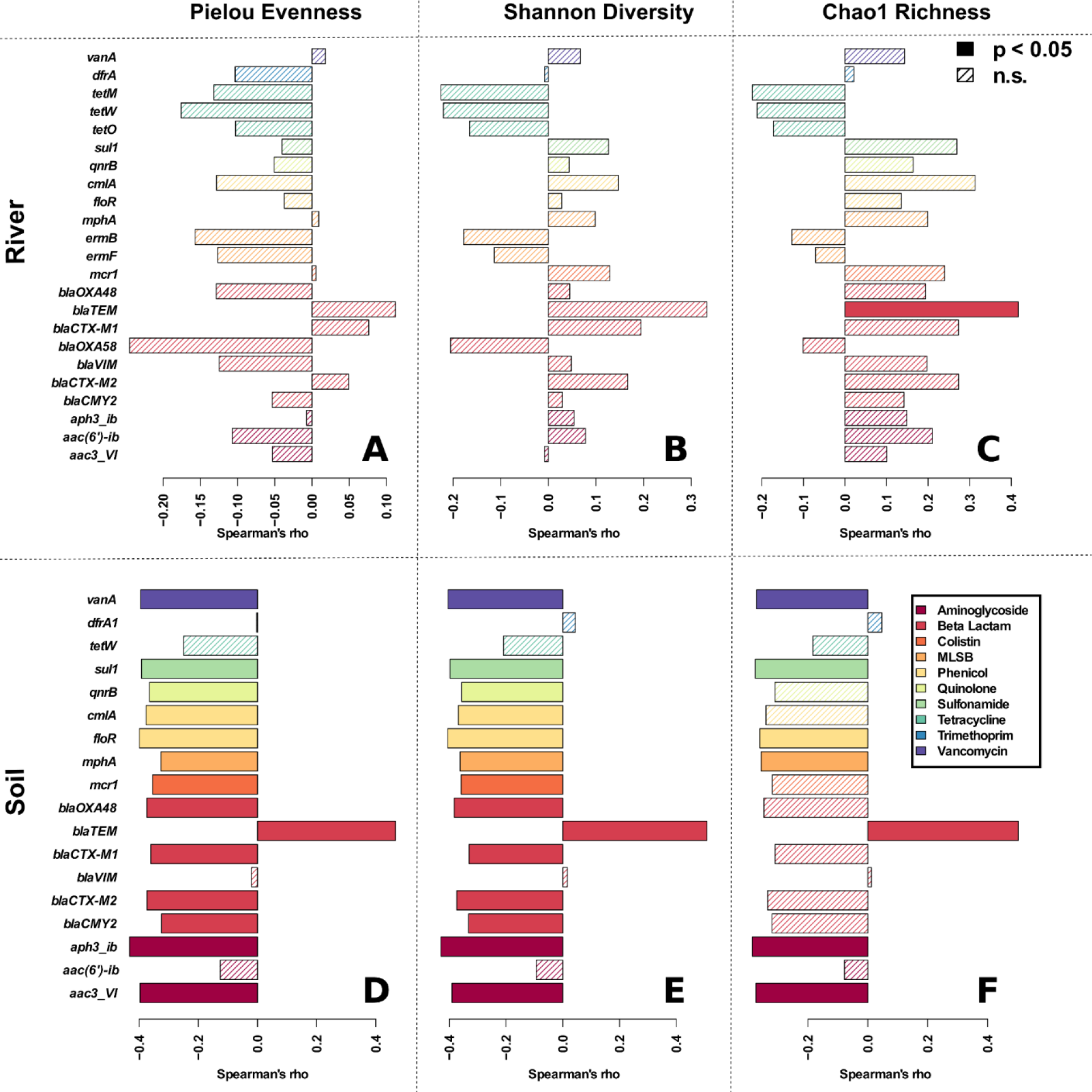
Correlation analysis of relative ARG abundance with observed diversity metrics based on Spearman rank correlation with Bonferroni correction for multiple testing. Correlations from river environmental samples with Pielou Evenness (A), Shannon Diversity (B) and Chao1 Richness (C). Correlations from soil environmental samples with Pielou Evenness (D), Shannon Diversity (E) and Chao1 Richness (F). Filled bars represent significant, while hatched bars represent non-significant correlations. Colors depict the class of antibiotic the ARG confers resistance to. Only ARGs that were detected in at least 25% of samples of a dataset were tested.

In contrast, a high number of significant negative correlations of relative ARG abundance with the different diversity indices were observed based on Spearman Rank correlation analysis in the soil dataset (Fig. 5 D-F). Pielou evenness and Shannon diversity displayed the most significant correlations with 13 of the 18 tested ARG relative abundances being negatively correlated, while six ARGs were negatively correlated with Chao1 richness. ARGs negatively correlated with diversity were widely distributed across antibiotic classes. Similar to the river environment, the *blaTEM* gene was the main exception from the observed trend and positively correlated to either diversity metric (RS= 0.47-0.51, all p < 0.05, Fig. 5 D-F). Despite this outlier, a general trend for the correlation between relative ARG abundance and diversity was observed for the soil dataset, with the average correlation coefficients of all tested ARGs being both negative and significantly different from 0 for Chao1 richness (RS = -0.257 ± 0.239, p = 0.0004), Pielou evenness (RS = -0.234 ± 0.228, p = 0.0001) and Shannon diversity (RS = -0.267 ± 0.223, p = 0.0003, Fig. 5 D-F). Finally, contrary to ARGs, none of the five indicator genes for MGEs tested (the class1 integron integrase gene *intI1*, the *IncP* plasmid *oriT*, the *IncW* plasmid *trwAB* gene, the *ORF37* gene of *IS26* and the Tn*5* transposase gene) displayed any correlation with either of the diversity indices in any of the datasets (all p>0.05).

## Discussion

Here we demonstrate based on analysis of a pan-European sampling campaign that communities of high bacterial diversity are more resistant to ARG pervasiveness, and that diversity might serve as a barrier to the long-term invasion and establishment of ARGs into environmental endemic microbiomes. Both, the number of detectable ARGs as well as a majority of the individual relative ARG abundances were negatively correlated with the diversity indices observed in soil. Among these indices, Pielou evenness was most significantly negatively correlated to the number of detected ARGs and their relative abundances. While this possible effect of community diversity on long-term invasion and establishment of resistant bacteria was highly visible and frequently statistically significant in the structured soil environment, it was barely observed in the more dynamic river environment.

Within the context of these results, the effects of diversity on invasion of AMR needs to be assessed individually for the successive steps that make up a successful invasion event, namely 1) introduction of the invader, 2) its establishment, 3) its growth and spread along with 4) its impact in the new microbial community [17]. The initial introduction of each invader is primarily of stochastic nature and does not rely on biological interactions with the indigenous community [17,20]. Consequently, the success of these initial introduction events depends on the quantity of invaders present, also known as the propagule pressure, together with the level of physical interaction of these invaders [21,44]. In this study, the samples originated from low impacted soils and rivers across Europe. Here, we define low impact as being not in direct proximity to the release of bacteria enriched in ARGs through anthropic action such as treated wastewater effluents [45,46] or manure [37,47]. The propagule pressure - the number of invaders harboring ARGs that were introduced into these environments - can be assumed low at the time of sampling and was likely low in the past. However, there is a high probability that bacteria with ARGs acquired in the antibiotic era occurred, nevertheless, at some rate (e.g. through human presence or transport by wild and domestic animals, including defecation, wet and dry atmospheric deposition). Consequently, it can be assumed that any increase in the resistomes in our samples are unlikely to stem from recent pollution events, but rather from past invasion events that manifest on top of the more or less universal background levels of resistance recently determined for a number of environments [48,49]. Increases in ARG occurrence and relative abundance would hence result from the accumulation of invasion success of previous repetitive introductions of invaders over time that have been able to establish themselves in the autochthonous microbiome or left their mobile ARG load behind, if we consider that bacteria from the human or animal spheres are regularly not fully fit to be long-term maintained in environmental microbiomes.

Contrary to the original introduction step of the invader, where biological interactions play only a minor role, the interactions with the local community are highly relevant during the subsequent establishment and growth phases. The internal resistance of the indigenous community towards invasion, e.g., its biotic barriers, have to be overcome to lead to the successful establishment of the invader [17,20] and can lead to the maintenance of ARGs in the community, when transient invasion success is long enough to allow for gene transfer [24]. Here we suggest that in the long term, microbial diversity might provide a biotic barrier, hindering ARG success of invasion in the low impacted soil microbial communities. However, no such effect could be observed for river communities. In the context of ARB such diversity effects have earlier been mainly demonstrated in short-term laboratory experiments for both types of environments using soil microcosms [26] and laboratory river flume experiments [27]. Still, in these experiments, diversity was artificially lowered to non-natural levels, which made it difficult to evaluate if such effects would equally be observable in the environment across natural biodiversity gradients in the long-term. In our analysis we make the implicit assumption that the present-day diversity is indicative also of past diversity, or at least of differences in diversity between sites and that both soil and riverbed microbiomes act as records of the long-term impacts endured by those microbial communities. This assumption appears likely to be correct in the case of forest soils, which are supposedly an environment typically stable over decades or more [50]. However, the assumption could be challenged regarding riverbeds, which are more likely exposed to considerably different conditions in the past (considering e.g., droughts, changes in water quality of European rivers over recent decades, etc.) [51]. This aspect may have contributed to the contrasting results in these environments.

The observed differences between the two datasets regarding the correlations between diversity and ARGs abundances likely originate from the different nature of the two environments. The resistance of the community to the invasion processes is directly related to the number of available niches for invader establishment, with more diverse communities providing less available niche spaces, referred to in macro-ecology as the “diversity-invasion effect” [20]. In the stationary and structured soil environment, the amount of available niches rarely changes and is, once occupied by a diverse community, able to reach a steady-state [52]. Under these circumstances it is unlikely that niches open up for invaders as no major loss of community members is to be expected. Within this context it is also unsurprising that community evenness is more strongly correlated with lower ARG abundance in soil than community richness. In highly even communities, a higher number of bacterial species are abundant enough to fully occupy their specific available niche space [53]. Rich communities have an intrinsically increased potential for their populations to occupy a higher number of different niches [54]. However, once a certain level of richness is reached, niche occupation moves towards saturation with additional species playing only a minor role [55]. Increased richness might then lead to increased competition and potential shifts within the metabolic networks, with novel niches becoming available to be exploited by the invading strains. With these two mechanisms at play, the negative correlation of richness with relative ARG abundances remains weaker than that observed for evenness or Shannon diversity.

In the river microbiomes, microbial diversity and available niches are potentially rather transient than long-term established due to the constant currents, biofilm adhesion and microbial dispersion events and alterations in nutrient availability [56]. Therefore, the effect of diversity on the long-term establishment of ARGs, which was observed for soil, may in the river microbiomes be masked by the dynamic nature of this environment. This difference between less (soil) and more (riverbed) dynamic environments is also displayed by the high number of ARGs being correlated in abundance in soil compared to the river dataset. While in soil the majority of abundant ARGs are present and affected by the described processes, in rivers ARGs rather appear to be more independent of each other based on dynamic processes.

Unlike for biotic invaders, such as alien bacteria, the invasion success of genetic invaders, such as ARGs, goes far beyond the successful establishment and growth of their hosts in the novel environment. A prolonged residence time of the host can favor the spread of mobile ARGs via horizontal gene transfer from the invading to the indigenous bacteria, thus leading to an even higher persistence of ARGs [24]. This can even be the case if the host’s invasion is only transiently successful and invaders are lost after some time due to high community resilience in the observed environment. Mobile ARGs encoded on plasmids are able to spread from an invading donor strain to highly diverse proportions of soil and water derived microbial communities, with bacteria belonging to over 25 different phyla able to receive individual resistance encoding plasmids [57–61]. However, effects of community diversity on the efficiency of horizontal gene transfer and the maintenance of plasmids in the community consist of a complex interplay of different mechanisms and remain difficult to predict. On the one hand, at higher diversity an increased number of potential plasmid hosts and conjugation partners are available that can lead to increased plasmid maintenance and transferability in the community [62,63] increasing the chance of transfer to a highly competitive host. On the other hand, in more diverse communities it can be harder to encounter a permissive conjugation partner, which reduces transferability due to this dilution effect [64]. Further, competition with other community members might increase the costs of resistance [65] and could ultimately drive the loss of ARG hosting plasmids from the community [66]. This loss process would be expected to be elevated in more diverse communities with better competitors. The complex interplay of mechanisms leading to a poor predictability of effects is equally represented in our soil dataset, where unlike for ARGs no clear correlation for MGEs (e.g., *intI1*, IncP plasmid *oriT*) with diversity could be established. However, a good indication that community diversity might also limit the horizontal acquisition of mobile ARGs from invading bacteria is that diversity is also negatively correlated with the number of detected ARGs in the soil dataset. This is according to ecological theory, where species diversity is not always immediately implying a higher degree of genetic diversity [67,68]. Still, assuming that, in the long-term, invaders harboring the tested ARGs reach each of the tested communities it becomes apparent that an increasing number of ARGs are not successfully retained in those communities of higher diversity. If this is due to a shorter residence time of the invader, the above discussed increased competition or dilution effects needs future research.

## Conclusions

In summary, we display that the microbial diversity within a given environment could affect the proliferation of AMR within and through this environment. Considering sites of low anthropogenic impact, the observed negative correlation of diversity with detection and abundance of ARGs in soils compared to river ecosystems can be directly connected to the intrinsic characteristics of the specific environments within the framework of invasion theory as well as horizontal gene transfer dynamics.

Natural environments, such as rivers and soils, may play a key role in AMR development and proliferation. The characteristics of the individual environment, its texture, its dynamism as well as the diversity of the resident microbial community could define its role as a source or a barrier to AMR dissemination. We present support that in the structured soil environment, a high bacterial diversity might indeed serve as a barrier to the long-term invasion and establishment of ARGs in the autochthonous microbiome. Our results point to a previously overlooked benefit of healthy environments, with diverse microbial communities, providing natural barrier effects to the proliferation of AMR, thus clearly displaying how environmental and human health are immediately interconnected through the One Health concept. Furthermore, such barrier effects can be exploited within soil ecosystem management, for example, in defining optimal locations for aquifer recharge through wastewater reuse. Here choosing locations with a high intrinsic diversity could be beneficial in limiting the spread of wastewater born ARGs. To achieve this, the role of microbial diversity in the dissemination and mobilization of AMR markers requires a closer look through targeted experiments aimed at elucidating the exact mechanisms that limit the proliferation of resistance determinants and how exploiting such natural barrier effects could have cascading effects on the ecosystem biodiversity.

## Declarations

### Ethics approval and consent to participate

Not applicable

### Consent for publication

Not applicable

### Availability of data and material

The datasets supporting the conclusions of this article are included within the article and its additional files. Original sequencing data is available in the NCBI sequencing read archive under project accession number PRJNA948643 (https://www.ncbi.nlm.nih.gov/bioproject/PRJNA948643), with individual sample identifiers given in tables S1 & S2.

## Author contributions

Conceptualization of the study and sampling strategy: UK, GG, EC, XB, SG, AGS, ES, CC, NK, MP, FW, MW, HB, CM, TUB; Identification of national sampling locations, sampling, metadata collection & sample processing: UK, GG, EC, XB, ID, SG, AGS, UO, ER, MS, ES, AT-M, CC, NK, MP, JV, FW, MW, HB, CM, TUB; Sampling coordination, sequencing and HT-qPCR data acquisition: UK, TUB; 16S sequence analysis: GG, EC; HT-qPCR analysis: UK; Data curation and validation: UK, GG, EC; Correlation and network analysis: UK, GG, EC, DK, PF; Data interpretation: UK, GG, EC, HB, CM, TUB; Visualization of data: UK, GG, EC, DK, PF; Funding acquisition: CC, NK, MP, FW, MW, HB, CM, TUB; Supervision: UK, CC, NK, MP, JV, FW, MW, HB, CM, TUB; Writing - original draft: UK, GG, EC; Writing - review and editing: all authors. All authors have read and approved the final version of the manuscript.

## Acknowledgements

We give a special thanks to Rosi Siber for creating the GIS Map (Figure S1). We thank all the local experts on soils and rivers in the different countries for advice regarding sample location identification.

