## Supplementary Information for "Microbiome diversity: A barrier to the environmental spread of antimicrobial resistance?"

* Contributed equally to this work

[#] Corresponding author

Corresponding author address:

Technische Universität Dresden

Institute of Hydrobiology

Zellescher Weg 40

01217 Dresden

Germany

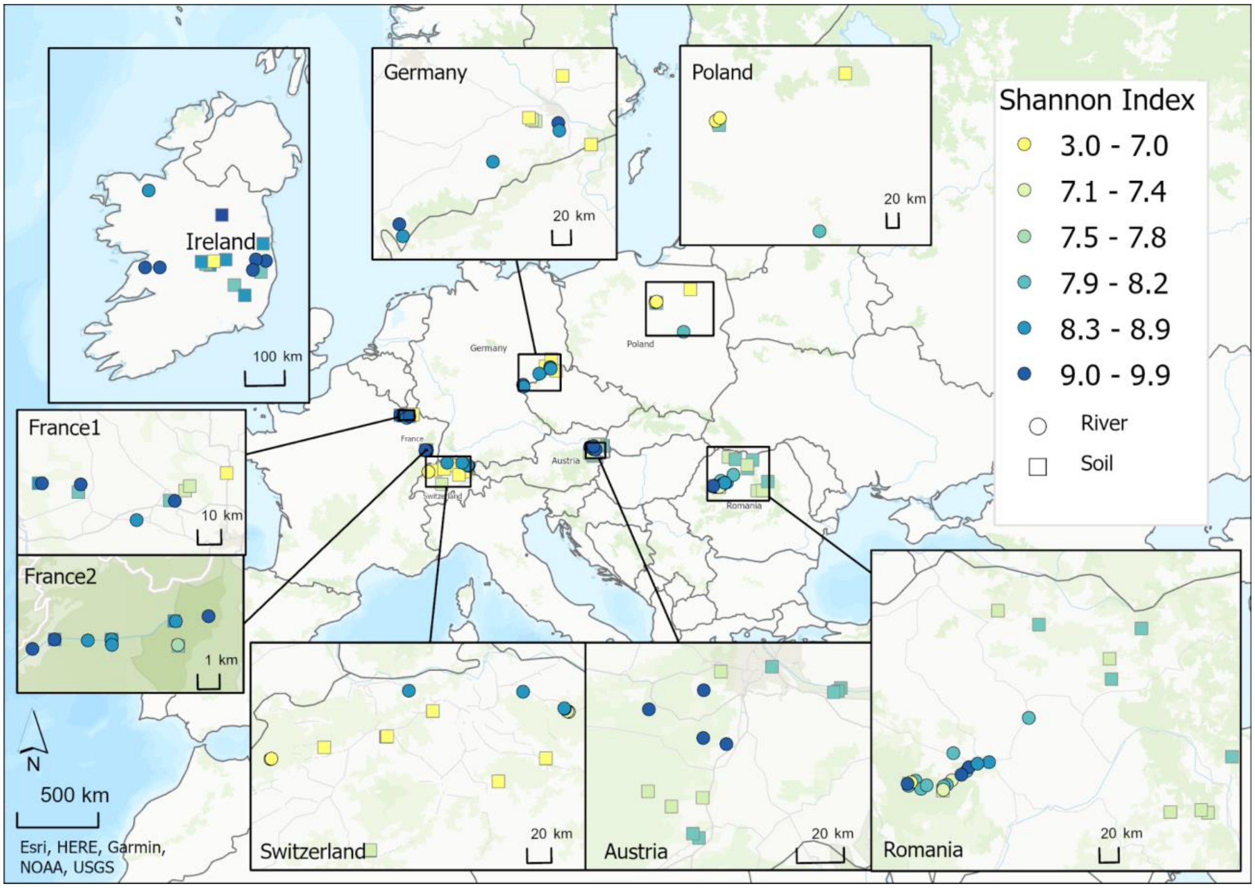


**Figure S1. Sampling locations across seven European countries**

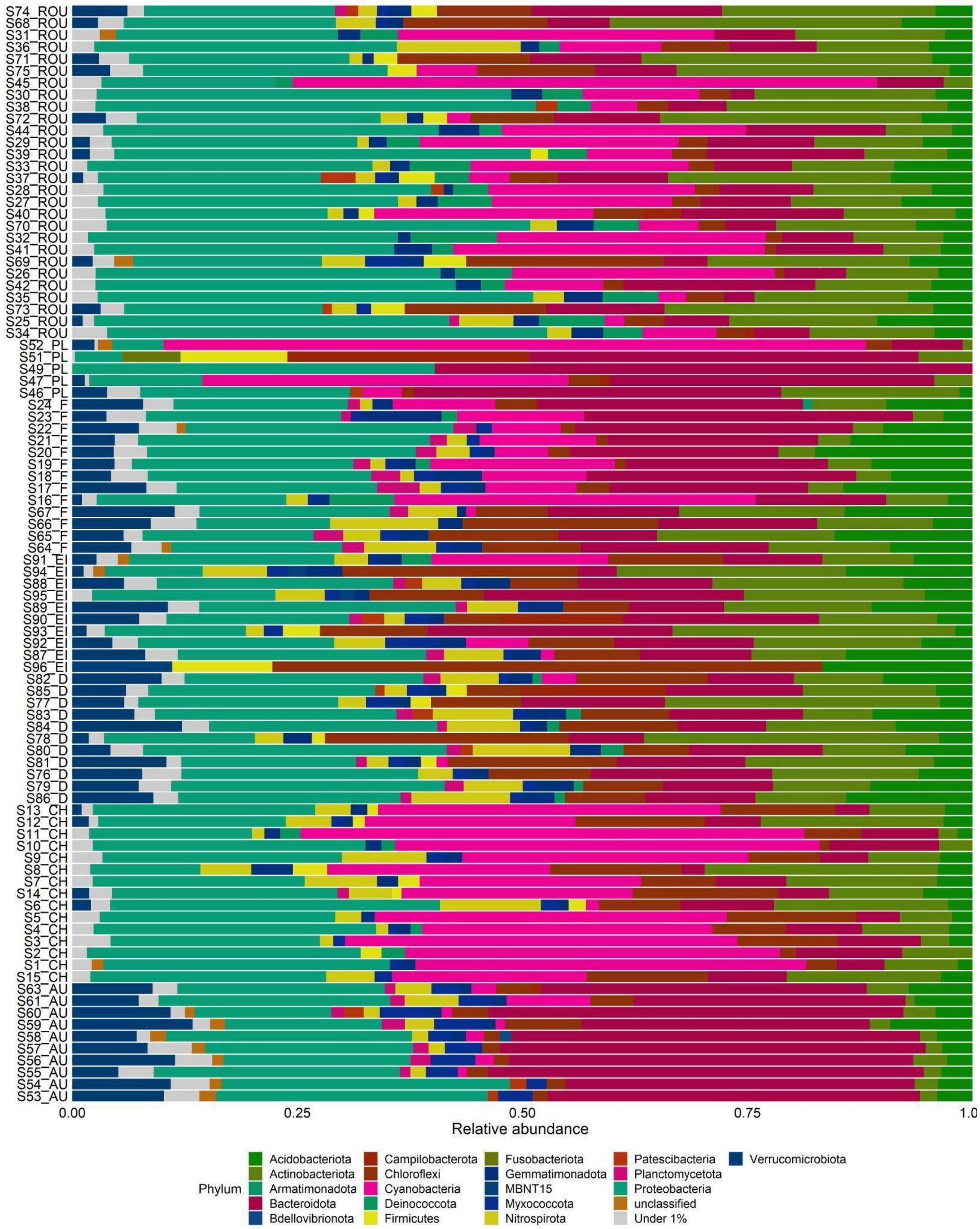


**Figure S2. Barplot of phylum diversity in all rivers sampled across 7 countries. For each sample, the phyla with less than 1% relative abundance were clustered as “Under 1%” to help with visualization.**

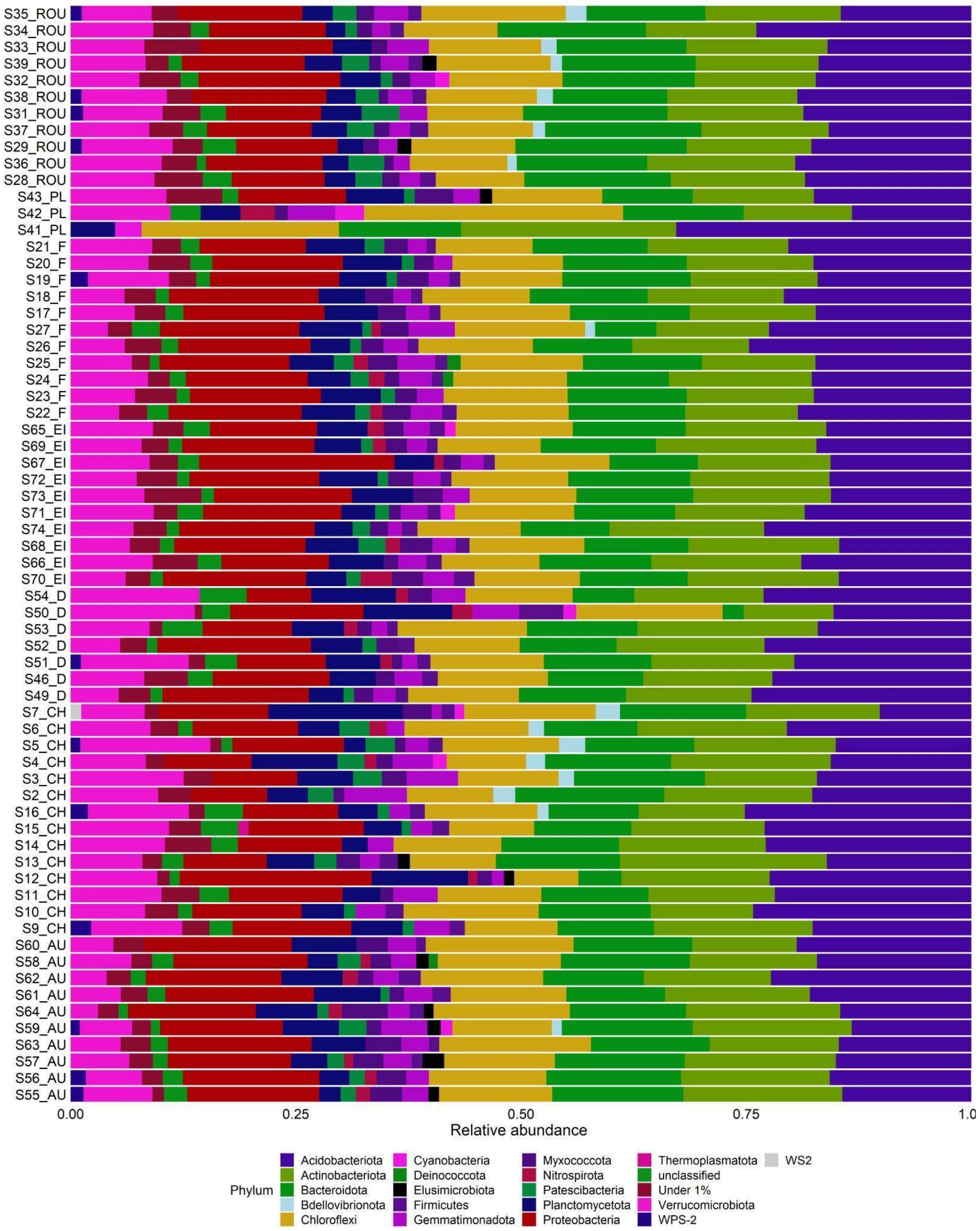


**Figure S3. Barplot of phylum diversity in all soil sampled across 7 countries. For each sample, the phyla with less than 1% relative abundance were clustered as "Under 1%" to help with visualization.**

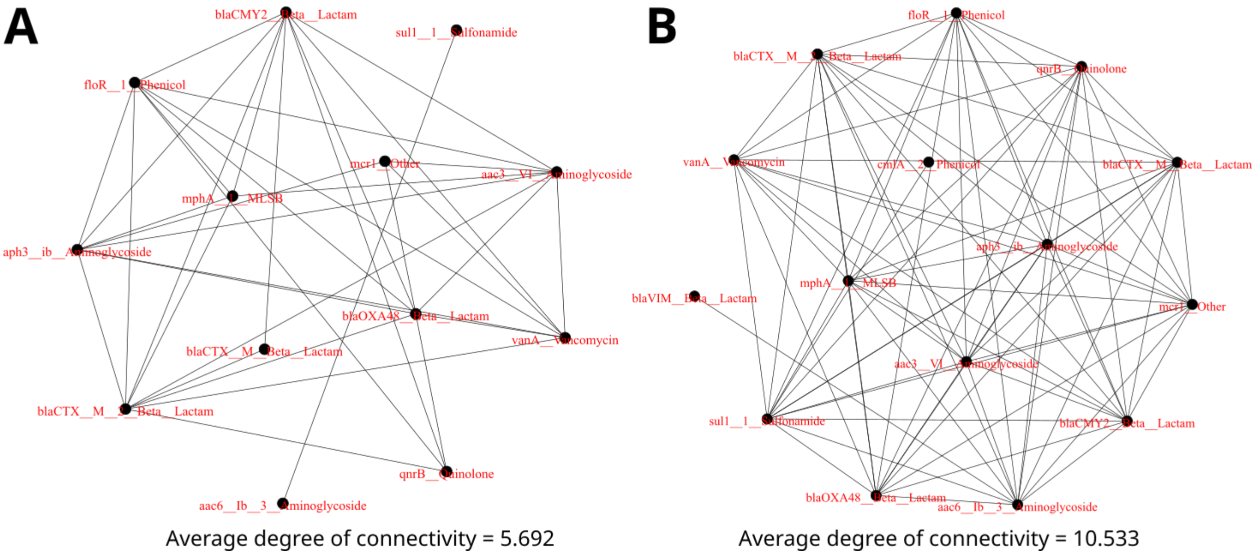


**Figure S4: Network analysis displaying the connectivity in relative abundance between the different ARGs in river (A) and soil samples (B) based on significant ($p < 0.05$) Spearman correlations with correlation coefficients of $|\rho| > 0.75$.**
